# Evolutionary inference from *Q*_ST_-*F*_ST_ comparisons: disentangling local adaptation from altitudinal gradient selection in snapdragon plants

**DOI:** 10.1101/385377

**Authors:** Sara Marin, Juliette Archambeau, Vincent Bonhomme, Mylène Lascoste, Benoit Pujol

**Affiliations:** Laboratoire Évolution & Diversité Biologique (EDB UMR 5174), Université Fédérale de Toulouse Midi-Pyrénées, CNRS, IRD, UPS, Toulouse, France; BIOGECO, INRA, University of Bordeaux, Pessac, France; Institut des Sciences de l’Évolution (ISEM), équipe “Dynamique de la biodiversité, anthropo-écologie”, UMR 5554, Université de Montpellier, CNRS, IRD, EPHE, Place Eugène Bataillon, Cc 065, 34095 Montpellier cedex 05, France; EPHE, PSL Research University, UPVD, CNRS, USR 3278 CRIOBE, F-66360 Perpignan, France.

**Keywords:** Local adaptation, altitudinal gradient, climate change, quantitative genetics, *Antirrhinum majus*

## Abstract

Phenotypic differentiation among natural populations can be explained by natural selection or by neutral processes such as drift. There are many examples in the literature where comparing the effects of these processes on multiple populations has allowed the detection of local adaptation. However, these studies rarely identify the agents of selection. Whether population adaptive divergence is caused by local features of the environment, or by the environmental demand emerging at a more global scale, for example along altitudinal gradients, is a question that remains poorly investigated. Here, we measured neutral genetic (*F*_ST_) and quantitative genetic (*Q*_ST_) differentiation among 13 populations of snapdragon plants (*Antirrhinum majus*) in a common garden experiment. We found low but significant genetic differentiation at putatively neutral markers, which supports the hypothesis of either ongoing pervasive homogenisation via gene flow between diverged populations or reproductive isolation between disconnected populations. Our results also support the hypothesis of local adaptation involving phenological, morphological, reproductive and functional traits. They also showed that phenotypic differentiation increased with altitude for traits reflecting the reproduction and the phenology of plants, thereby confirming the role of such traits in their adaptation to environmental differences associated with altitude. Our approach allowed us to identify candidate traits for the adaptation to climate change in snapdragon plants. Our findings imply that environmental conditions changing with altitude, such as the climatic envelope, influenced the adaptation of multiple populations of snapdragon plants on the top of their adaptation to local environmental features. They also have implications for the study of adaptive evolution in structured populations because they highlight the need to disentangle the adaptation of plant populations to climate envelopes and altitude from the confounding effects of selective pressures acting specifically at the local scale of a population.

## INTRODUCTION

Local adaptation - the evolutionary response to local selection that makes populations fitter in their own local habitat than in any other population’s local habitat - is widespread in both plant and animal species (Kawecki & Ebert, 2004; Tuomas Leinonen, McCairns, O’Hara, & Merila, 2013). One very popular indirect approach to assess whether local adaptation has shaped trait quantitative genetic variation is the *Q*_ST_-*F*_ST_ comparison (McKay & Latta, 2002; Merila & Crnokrak, 2001; Spitze, 1993). It consists in the comparison of population genetic differentiation (e.g., D and *F*_ST_) estimated for putatively neutral molecular markers (Edelaar & Björklund, 2011; Rousset, 1997) with population quantitative genetic differentiation (*Q*_ST_) estimated for phenotypic traits of interest (Whitlock, 2008). The objective is to test whether trait quantitative genetic differentiation among populations is more likely the result of divergent selection or neutral evolutionary divergence (e.g., as a result of drift). This approach has three main advantages. First, the necessary data is relatively easy to acquire (phenotypic measures and genotypes). Second, population genetic differentiation statistics are relatively easy to estimate. Third, conclusions are mostly derived straightforwardly from results: *Q*_ST_>*F*_ST_ is taken as evidence for local adaptation, *Q*_ST_<*F*_ST_ is taken as evidence for stabilizing selection, and *Q*_ST_=*F*_ST_ means that selection is not required to explain population differentiation because neutral evolution might be responsible for the divergence. There has been some debate around the accuracy of *Q*_ST_-*F*_ST_ approaches. As a result, the advantages and limits of this method are relatively well known, and a variety of possible methodological adjustments to the estimation of differentiation are available (De Villemereuil & Gaggiotti, 2015; Edelaar, Burraco, & Gomez-Mestre, 2011; Ovaskainen, Karhunen, Zheng, Arias, & Merilä, 2011; Whitlock, 2008; Whitlock & Gilbert, 2012). Although some of these adjustments were later discarded (Edelaar et al., 2011; Whitlock, 2011), more than thirty years of extensive use have resulted in wide acknowledgement of the *Q*_ST_-*F*_ST_ approach as a method testing for the hypothesis that local adaptation might explain observed patterns of differentiation. The influence of environmental effects on phenotypic variation that might result in biasing *Q*_ST_-*F*_ST_ comparisons conducted directly in wild populations has also been debated (Pujol, Wilson, Ross, & Pannell, 2008). Yet, the inclusion of ecological factors in this type of analysis remains rare (Hangartner, Laurila, & Räsänen, 2011; Karhunen, Ovaskainen, Herczeg, & Merilä, 2014). Consequently, the identification of local habitat selective pressures that might drive local adaptation is often missing from *Q*_ST_-*F*_ST_ approaches.

The general absence of identified ecological drivers for local adaptation in *Q*_ST_-*F*_ST_ approaches is not surprising. This is likely because it is a correlative approach, which limits conclusions about causality, and because it is impossible to record and include in the analysis every ecological variable differing between populations. Consequently, confounding effects from unmeasured variables can never be ruled out. It is nevertheless possible to investigate the link between changes in population divergence and changes in ecological variables at the scale of multiple populations. This does not allow identifying the local features of the environment of every population that drive local adaptation *per se*. It can however shed light on traits that are involved with adaptation to a particular ecological background at a larger scale. In fact, the exploration of ecologically explicit adaptive scenarios based on *Q*_ST_-*F*_ST_ approaches that are not particularly “local” has broader implications in the actual context of environmental change (e.g., climate change, pollution) because they affect multiple populations simultaneously. For example, identifying candidate traits that are involved with adaptation to different climates has obvious implications for climate change remediation (De Villemereuil, Mouterde, Gaggiotti, & Till-Bottraud). Other approaches targeting the molecular variation of adaptive genes also allow us to investigate the same question (Ahuja, de Vos, Bones, & Hall, 2010; Hoffmann & Sgro, 2011), but most of them cover less populations and are more demanding in time and resources. Breaking down this accessibility barrier by using less demanding approaches has implications for the generalisation of findings. It is therefore only logical that the general challenge of *Q*_ST_-*F*_ST_ approaches is shifting progressively from detecting the signature of local adaptation to identifying the ecological drivers of selection (Hangartner et al., 2011; Whitlock, 2008).

There is a growing number of studies investigating whether population quantitative genetic divergence is shaped by environmental constraints (Gomez-Mestre, Tejedo, & Ashley, 2004; Palo et al., 2003; Raeymaekers, Van Houdt, Larmuseau, Geldof, & Volckaert, 2007). For example, Hangartner et al. (2011) showed that population adaptive divergence in metamorphic size was associated with pond acidity in moor frogs. They also identified that other traits such as larval period and growth rate were involved with adaptation to latitude and predation. This was possible because they investigated whether the increase of environmental differences between populations was associated with an increase in *Q*_ST_ for various traits. This type of approach can ultimately inform us about the vulnerability of populations to climate change because comparing their adaptive divergence along altitudinal gradients can serve as a first-order approximation of the possible effects of ongoing climate change (Byars, Papst, & Hoffmann, 2007; Gonzalo-Turpin & Hazard, 2009). There is evidence for the adaptive signature of past climate change in multiple populations growing number of studies on this topic (De Villemereuil et al.; Luquet, Léna, Miaud, & Plénet, 2014; McKay et al., 2001; Muir, Biek, Thoma, & Mable, 2014). For example, Muir et al. (2014) showed that local adaptation had shaped the five traits they had measured in common frog populations by using a common garden experiment, and that two of these traits, i.e., larval period and growth rate, played a role in their adaptation to altitude. In plants, reciprocal transplants between altitudes that are used to compare plant fitness directly in their native habitat and in the foreign habitat have been mostly preferred to *Q*_ST_-*F*_ST_ indirect approaches that are conducted in common gardens (Angert, Schemske, & Geber, 2005; Etterson, 2004; Kim & Donohue, 2013). Using an indirect approach in controlled conditions can however present an advantage when the conditions for the reciprocal transplant cannot be met, e.g., when gene flow between populations on multiple sites that would result in mixing up the genomic background of all these populations cannot be avoided. In such cases, *Q*_ST_-*F*_ST_ comparisons along altitudinal gradients might present a rare opportunity for exploring hypotheses about candidate traits for the adaptation of plants to climatic differences.

Functional ecology studies predict that eco-physiological and reproductive traits play a crucial role in the adaptation of plants to the later and shorter availability of favourable environmental conditions that characterise seasons at higher altitudes (Körner, 1999). Selection should therefore favour smaller plants, germinating later but reproducing earlier, with faster growth at higher altitude. By reciprocity, plants from lower altitude that are confronted with warmer climates should present opposite adaptations. However, these hypotheses have rarely been tested using empirical data from multiple natural populations across altitudinal gradients. Here we present the results from a common garden experiment where we evaluated the adaptive significance of multiple phenological, morphological and functional traits in populations of snapdragon plants (*Antirrhinum majus* L.). We estimated neutral genetic differentiation (*F*_ST_), trait heritability (*h*²), and quantitative genetic differentiation (*Q*_ST_) among 13 *A. majus* populations. We estimated the genetic differentiation between populations at putatively neutral microsatellite markers (Debout, Lhuillier, Malé, Pujol, & Thébaud, 2012; Pujol et al., 2017) to test for the hypothesis that gene flow was limited between populations, which sets the stage for local adaptation. We then tested for the hypothesis of local adaptation by comparing *Q*_ST_ and *F*_ST_. Finally, we investigated whether quantitative genetic differentiation increased with altitude, with the hypothesis that broader environmental changes associated with altitude, such as the climatic envelope, can explain more population adaptive trait differentiation than local population differences. Our approach ultimately participates to the collective evaluation of the *Q*_ST_-*F*_ST_ approach, which is generally used for the study of local adaptation in structured populations, as a tool to identify the adaptive response of local populations to altitudinal gradients.

## MATERIAL AND METHODS

### Study system

*Antirrhinum majus* L. (Plantaginaceae) is a hermaphroditic, self-incompatible, short-lived perennial, which produces annual inflorescences with zygomorphic flowers. These flowers are arranged in raceme and produced at the end of the stem, thereby preventing further vegetative elongation (terminal flowering). This plant is characterized by a patchy distribution in southern Europe centred over the Pyrenees Mountains. Two subspecies, that are interfertile, occupy this geographic area: *A. m.* ssp. *pseudomajus*, characterized by magenta flowers, and *A. m.* ssp. *striatum*, characterized by yellow flowers. The geographic range of *A. m. striatum* is surrounded by the range of *A. m. pseudomajus* (Khimoun et al., 2011). Within the species geographic range, the two subspecies are distributed parapatrically and come into contact at the margins of their ranges (Andalo et al., 2010). The transition between subspecies in the contact zones occurs over a very short distance (<1km) (Whibley et al., 2006). In the west part and the east part of the contact perimeter, there is evidence for introgressive hybridization between *A. m.* ssp. *pseudomajus* and *A. m.* ssp. *striatum* (Khimoun et al., 2011). The clear separation between the distribution of *A. m.* ssp. *pseudomajus* and *A. m.* ssp. *striatum* is not explained by habitat differences, as illustrated by the substantial overlap of environmental conditions between the two species (Khimoun et al., 2013). In fact, the range of environmental conditions characterizing the ecological niche of *A. m.* ssp. *pseudomajus* is almost twice as large as that of *A. m.* ssp. *striatum* (Khimoun et al., 2013). Both subspecies occur from sea level to an altitude of 1900 m (Andalo et al., 2010), on limestone or siliceous substrates and in habitats with contrasted moisture regimes (rainfall 500-1000 mm per year), where they form restricted patches mostly in rocky outcrops and screes. *A. majus* thrives in disturbed habitats, and is especially common along roadside and railway embankments. Thirteen wild populations were sampled in 2011 across the geographic range (between north-eastern Spain and south-western France), with four populations of *A. m.* ssp. *striatum*, six populations of *A. m.* ssp. *pseudomajus*, and three populations (one *A. m.* ssp. *striatum*, and two *A. m.* ssp. *pseudomajus*) from the contact zones (Figure 1; Table S1, Supporting Information). For each subspecies and in the contact zone, we sampled populations from low and high altitude habitats. This is because populations sampled along elevation gradients are likely to be confronted to contrasted environmental conditions (Figure S1, Supporting Information). Populations were sampled in different valleys or on different summits in order to avoid spatial autocorrelation in the data, and to avoid shared phylogeographic history between populations from similar altitudes.

**FIGURE 1.**
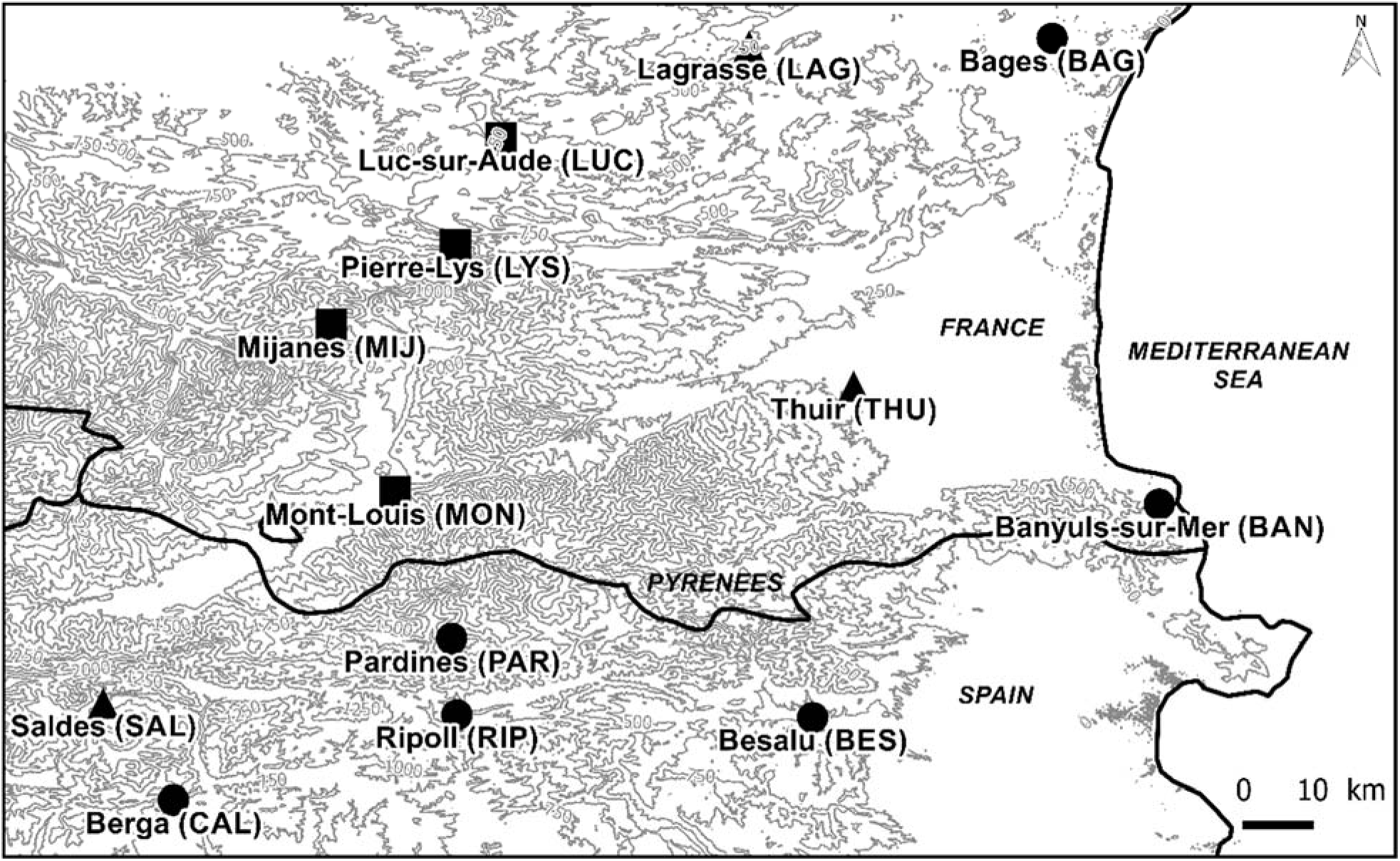
Map of *Antirrhinum majus* populations that were sampled across the geographic range of the species in Southern France. Dots represent *A. m.* ssp. *pseudomajus* populations, squares represent *A. m.* ssp. *striatum* populations and triangles represent populations from the contact zone. Grey lines represent elevation isoclines every 250 m.

### Common garden experiment

In each wild population, mature plants of *A. majus* were randomly sampled for seeds across their entire spatial distribution in October 2011 (Table S1, Supporting Information). We avoided maternal environmental effects in our common garden experiment by producing a seed bank from plants grown in similar conditions in a greenhouse at the CNRS Station of Experimental Ecology in Moulis. Seeds sampled in the wild were sown in spring 2012 in individual pots (9 × 9 × 10 cm) filled with universal compost. Plants germinated and grew with no nutrient addition under an average temperature from 15 to 28°C and weekly watering.

Mature plants were hand-pollinated during summer 2012. Crosses were conducted within populations where mates were assigned randomly. Seeds were stored in the seed bank (room temperature, dark, dry conditions). We sowed seeds from the seed bank in spring 2014 in a common garden at ENSFEA (Toulouse, France). Plants were grown outdoor in individual pots (9 × 9 × 10 cm) that were positioned in plastic containers (600 × 400 × 120 mm). Their growth was monitored during the summer 2014. Plants grew in pots filled with universal compost, with no nutrient addition, under outdoor climatic conditions (average month temperatures ranging from 20.6 to 21.5°C and cumulative monthly rainfall ranging from 28.3 to 73.4 mm). The bottom of each container was covered with an irrigation sheet (400 g.m-²) that allowed the regulation of compost moisture. Plants were supplied with water in case of prolonged drought. Damage caused by herbivorous insects were contained by using a wintering veil. This veil also limited pollination. The spatial location of each plant in the experimental setting was distributed randomly to avoid spatial effects on plants caused by uncontrolled environmental conditions.

### Molecular and phenotypic data

In terms of molecular markers, we genotyped the 13 different populations (*N* = 637 individuals) for 23 putatively neutral microsatellite markers that were developed for population genetic studies (Debout et al., 2012; Pujol et al., 2017). In terms of phenotypic traits, we measured multiple phenological, morphological, functional and reproductive characters on each individual. Phenological traits included germination date, flowering date, and time to flowering (the number of days separating the germination of a plant and the emergence of its first flower). At the time of first flowering, we measured vegetative traits: main stem basal diameter (Diameter), main stem vegetative length (Vegetative Length), number of vegetative nodes on the main stem (Nodes), total number of leaves on the plant (Leaves), and number of branches on the plant (Branches). We also measured these vegetative traits for plants that did not flower but stopped growing at the end of experiment. As a result, vegetative traits were characterised for all plants. At the end of the experiment, we also measured reproductive traits on flowering plants (*N* = 380): stem total length (Total Length), inflorescence length, number of flowers (Flowers). Additionally, we measured a functional trait when possible: specific leaf area (SLA). SLA was calculated as the ratio between the cumulated area of five mature but non senescent fresh leaves and their oven-dried mass (Pérez-Harguindeguy et al., 2016; Pujol, Salager, Beltran, Bousquet, & McKey, 2008). Leaf area was measured by using the R package Momocs v. 1.2.9 (Bonhomme, Picq, Gaucherel, & Claude, 2014). We also measured a developmental trait, average Internode Length for the total stem vegetative length.

### *Q*_ST_-*F*_ST_ comparison to evaluate local adaptation and altitudinal selection

To explore the effect of local adaptation on traits, we tested for the difference between the *Q*_ST_ observed value (global population quantitative genetic differentiation for a trait) and its expected value under the hypothesis of neutral evolution by using the approach of Whitlock and Guillaume (2009). As a result, we did not compared *Q*_ST_ with average *F*_ST_ but with the *Q*_ST_ expected distribution under a scenario of neutral evolution. This distribution of neutral *Q*_ST_ was obtained by using simulations parametrized on the basis of observed mean *F*_ST_ (neutral genetic differentiation at molecular markers) and predicted from the χ² distribution. The expected neutral *Q*_ST_ distribution for each trait was generated by bootstrapping (N = 10^5^). We then inferred *P* values associated with our point estimates of *Q*_ST_-*F*_ST_ differences. Finally, we used the modification by Lind et al. (2010) of the approach of Whitlock and Guillaume (2009) to estimate variance components. The choice of this method was motivated by the fact that it suits perfectly data like ours (population count around ten around ten, and availability of molecular markers such as microsatellites rather than genomic markers). All procedures were implemented in the R environment v. 3.5.0 (R-Core-Team, 2018). We then evaluated the correlation between matrices of pairwise population altitudinal and *F*_ST_ distances, and between pairwise population altitudinal and *Q*_ST_ distances. The significance (α=0.05) of these correlations was tested by using Mantel partial correlation tests (Mantel, 1967). Some drawbacks have been reported for this method, in particular in the presence of spatial autocorrelation. However, we limited the possibility for data spatial autocorrelation in altitude at the scale of populations by using a specific sampling scheme (see above). The option of using a Bayesian approach to overcome this issue was not available to us as it would have required using around 1000 molecular markers (Bradburd, Ralph, & Coop, 2013). We tested for Mantel partial correlations by using 9999 permutations (*mantel.rtest* command in the R package ade4 (Chessel, Dufour, & Thioulouse, 2004).

### Calculation of *F*_ST_, *h*² and *Q*_ST_

To compute *F*_ST_, we used estimates from the study by Pujol et al. (2017) for both pairwise population *F*_ST_ estimates and global population overall estimates. Potential effects on population genetic differentiation of geographic coordinates (Latitude and Longitude), pairwise geographic distance, subspecies and altitude were controlled for by using an AMOVA approach. Genetic diversity at each locus, which measure was required for the *Q*_ST_-*F*_ST_ comparative approach that we used, was estimated by using the GENETIX 4.05 program (Belkhir, Borsa, Chikhi, Raufaste, & Bonhomme, 2004). Narrow-sense heritabilities (*h*²) were estimated for each phenotypic trait. They were calculated as *h*² = *V*_A_ / *V*_P_, with *V*_P_ the population phenotypic variance and *V*_A_ the additive genetic variance. *V*_A_ was estimated as 2*V*_W_, with *V*_W_ being the within-population genetic variance. *V*_W_ was estimated by the among-family variance. We multiplied *V*_W_ by two in the calculation of *h*² because we used a full-sib crossing design (Roff, 1997). No environmental source of phenotypic variance due to the ecological conditions of the location of origin of populations could in theory bias *Q*_ST_ estimates because data was obtained from a common garden experiment (Benoit Pujol et al., 2008). Furthermore, at the scale of the common garden, plant locations were assigned randomly. *Q*_ST_ estimates were calculated by using the following formula (Spitze, 1993): *Q*_ST_ = *V*_B_ / (*V*_B_ + 2*V*_W_) with *V*_B_ being the trait genetic variance among populations. All variance components were estimated by using a Linear Mixed Model approach implemented in the R package lme4 v. 1.1.17 (Bates, 2005). When a variance component was not significant, it was considered as null in further calculations. In these models, population and family were included as random factors. We verified the normality of the distribution of residuals (Shapiro test) and their homoscedasticity (Breusch-Pagan test). When necessary (as for pairwise *Q*_ST_ calculation), data was linearized by using a square root transformation.

## RESULTS

### Local adaptation

Average phenotypic values and their standard errors were compared among the 13 experimental populations for the 13 analysed traits (Table S2, Supporting Information). For phenological traits, average germination dates of populations spanned across seven days (ranging from the 194^th^ to the 201^th^ calendar day of 2014) while the average flowering date spanned across 15 days (ranging from the 236^th^ to 251^th^ calendar day of 2014). Among populations, the time to flowering ranged from 43 to 59 days. For vegetative traits, amongst populations, Diameter ranged from 3.37 to 4.14 mm, Vegetative Length ranged from 19.95 to 35.74 cm, Nodes ranged from 10 to 17, Leaves ranged from 95 to 152, and Branches ranged from 12 to 20. Total Length, which included the length of the inflorescence, ranged from 25.18 to 54.49 cm. For reproductive traits, inflorescence length ranged from 6.86 to 20 cm and Flowers ranged from 6 to 19. Finally, we measured an average SLA ranging from 161.24 to 194.48 cm².g^-1^ and an Internode Length ranging from 1.75 to 2.43 cm.

Although not directly confronted to trait *Q*_ST_ values, we have represented on Figure 2 the low but significant average *F*_ST_ among populations (*F*_ST_ = 0.107, *P* < 0.001), which ranged from 5 to 20% in order to illustrate population neutral genetic differentiation (for more details on population pairwise neutral genetic differentiation, see Pujol et al., 2017). We found that most trait *Q*_ST_ values were not in the tail of the expected probability distribution of neutral traits, which means that the hypothesis of neutrality was rejected for these quantitative traits. Trait *Q*_ST_ values are represented on Figure 2. These values were generally significantly higher than the simulated distribution of their expected neutral values (Figure 2, Table S3, Supporting Information). However, for germination date, *Q*_ST_ was in fact lower than could be expected under neutrality. For time to flowering, *Q*_ST_ was in the tail of the expected probability distribution of neutral traits. Heritability estimates were generally comprised between 0.07 and 0.23. However, germination date, time to flowering, Nodes and Internode Length, had a heritability between 0.36 and 0.47 (Table S3, Supporting Information). Caution must be taken when considering *h*² values because its estimation can be biased by the estimation of *V*_A_. Indeed, *h*² was calculated on the basis of all the families, without taking into account the differences of *h*² between different populations. The family variance component was not significant and thereby considered null (*V*_W_ = 0) for the Leaves, making simulations by the approach of Whitlock and Guillaume (2009) impossible.

**FIGURE 2.**
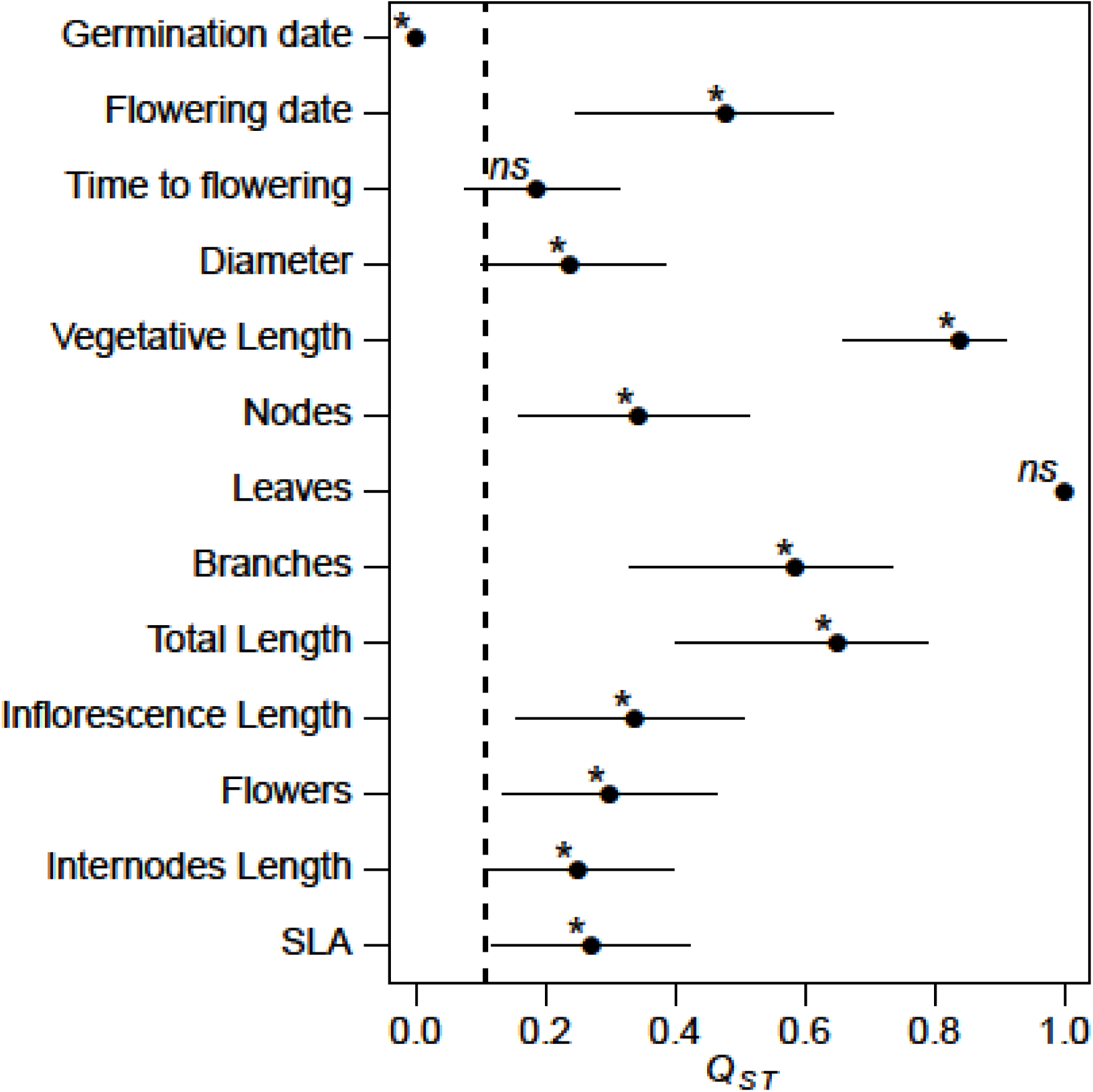
Values for observed *Q*_ST_ (represented by dots with its 95% confident interval indicated by black lines). *P*-values of statistical differences between observed values and the expected distribution of neutral traits are indicated by * for significant differences, or *ns* for non-significant differences. Average population *F*_ST_ is represented by the dashed line (*P* < 0.001).

### Increased quantitative genetic differentiation with altitude

The larger the difference in altitude was for a pair of populations, the higher was their neutral genetic differentiation, as estimated by *F*_ST_. This altitudinal effect was weak but significant (Figure 3). The relationship with altitude was also significant for quantitative genetic differentiation as measured by *Q*_ST_. It was only significant for three traits out of the 13 traits that we measured (germination date, inflorescence length, Flowers). The slope of the altitudinal increase in differentiation was one order of magnitude higher for *Q*_ST_ than *F*_ST_ (Figure 3). Indeed, we found that per unit elevation of 500 meters, *Q*_ST_ of germination date increased by 11%, *Q*_ST_ of inflorescence length increased by 12.5%, and *Q*_ST_ of Flowers increased by 13.5%, while *F*_ST_ increased by 1.5%.

**FIGURE 3.**
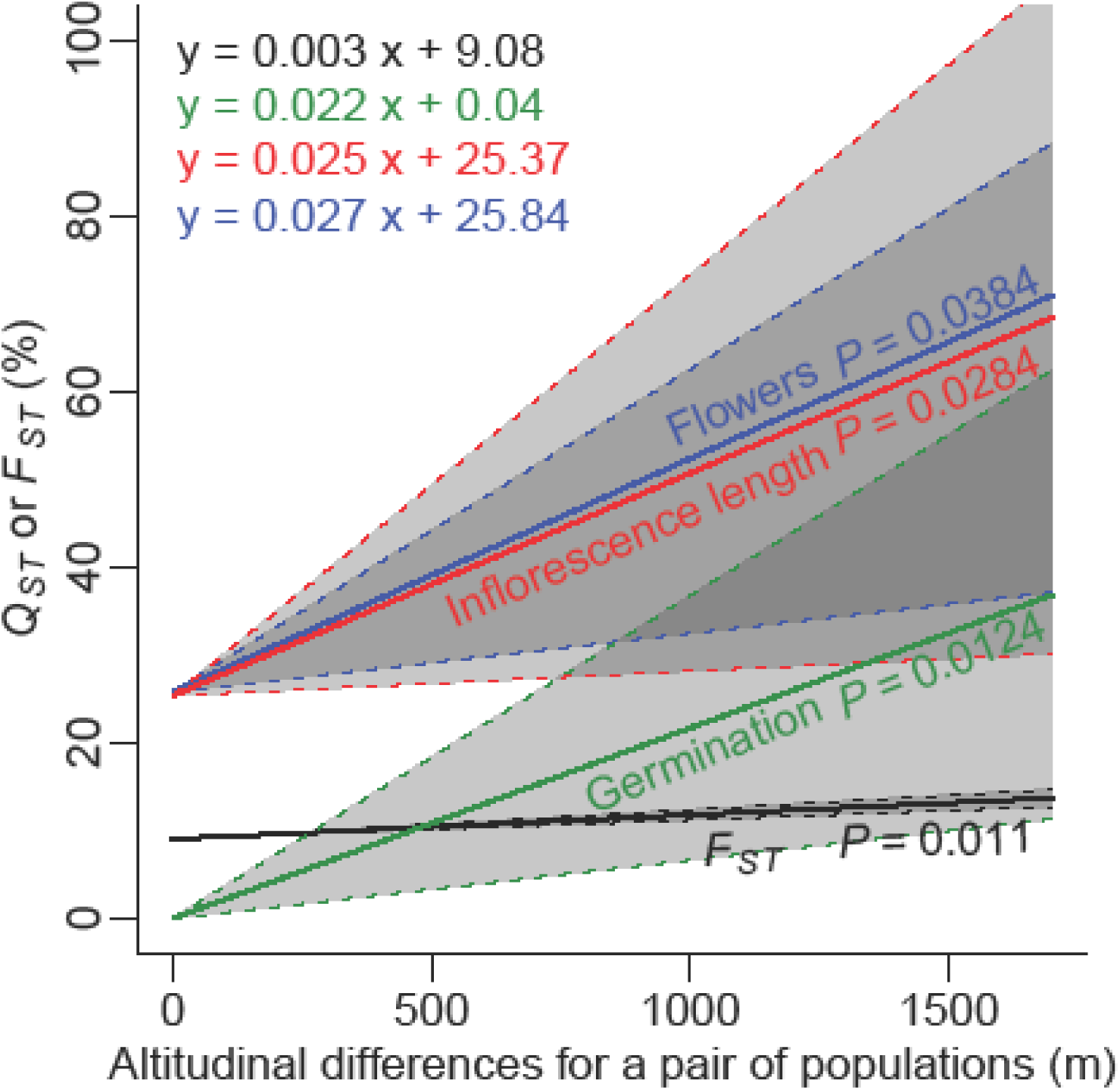
Linear regression (coloured lines) of population pairwise quantitative trait differentiation (*Q*_ST_) on population pairwise differences in altitude (m) for Germination date, Inflorescence length and Flowers. Linear regression (grey line) of the population pairwise neutral genetic differentiation (*F*_ST_) on population pairwise differences in altitude (m). Regression equations and levels of significance (*P*), obtained by using Mantel tests are presented in the figure. 95% confidence intervals are indicated by shaded areas.

### Changes in phenotypic values with altitude

Germination occurred significantly later in experimental populations that originated from higher altitudes. Germination date increased by 0.59% (1.15 days) per unit elevation of 500 meters (Figure 4a). Plants from populations originating from higher altitudes showed a significant reduction in the size of their inflorescence and in the number of flowers. Inflorescence length decreased by 13.14%, and Flowers decreased by 12.43% per unit elevation of 500 meters (Figure 4b, c).

**FIGURE 4.**
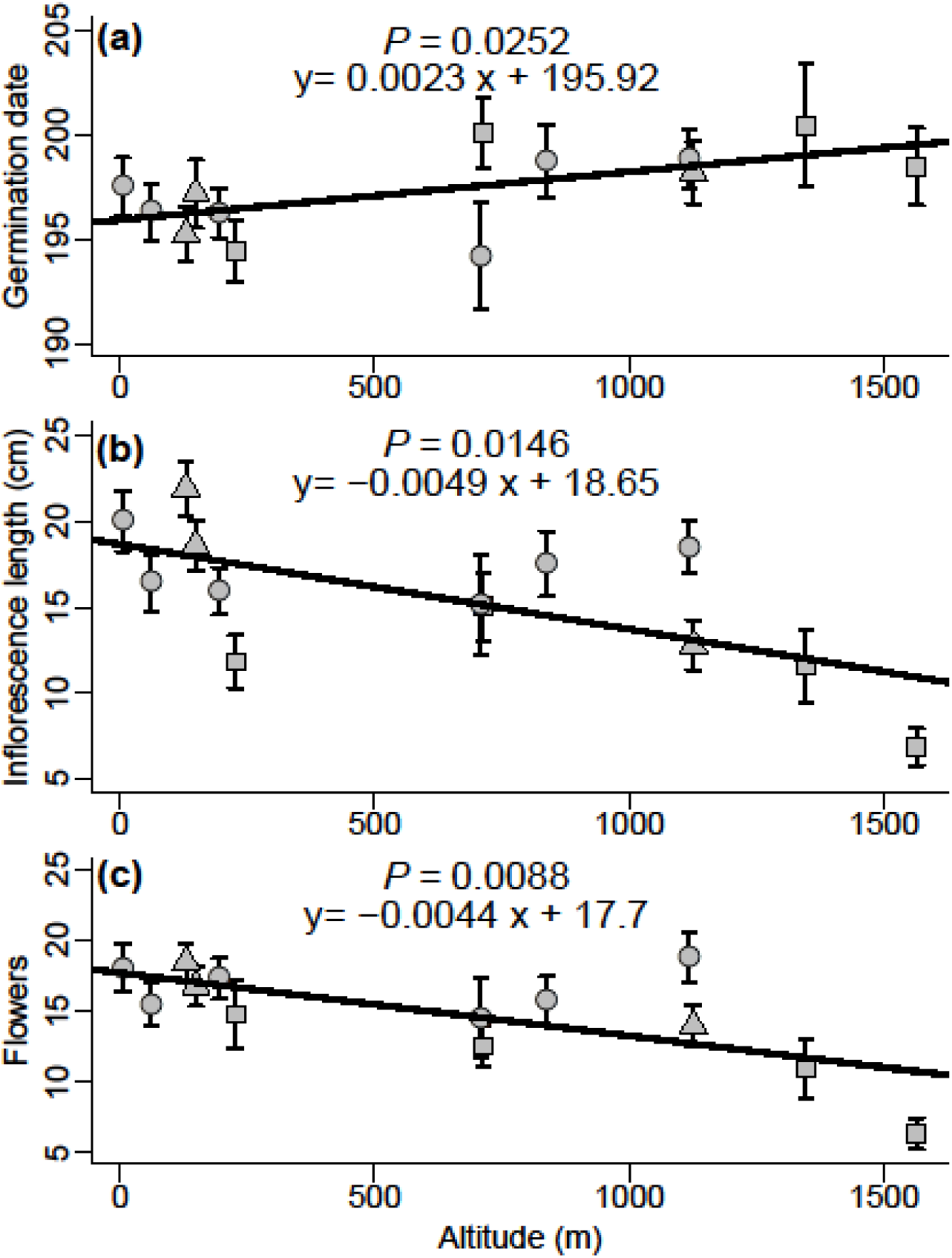
Variation of mean (±SE) per populations for Germination date (a), Inflorescence length (b) and Flowers (c) with altitude. Associated regression (solid line) equations and level of significance (*P*) are presented for each phenotypic trait. Dots represent *A. m.* ssp. *pseudomajus* populations, squares represent *A. m.* ssp. *striatum* populations and triangles represent populations from the contact zone.

## DISCUSSION

Our results showed that neutral evolution and local adaptation could not explain on their own the extent to which *A. majus* populations from the highest altitudes had diverged from lower populations. Phenotypic differentiation increased significantly with altitude for the germination date, inflorescence size and number of flowers, with more than a two-fold increase at the highest altitudes. Our results also provide support for the hypothesis that neutral evolution had the potential to shape the diversity of *A. majus* populations over their geographic range. However, this potential could only account for around 10% of population phenotypic differentiation. On the top of their neutral divergence, it was necessary to invoke local adaptation for explaining how *A. majus* populations diverged from each other at multiple phenological, morphological, reproductive and functional traits. Our findings comfort the emerging idea that *Q*_ST_-*F*_ST_ comparisons can be used to go further than detecting the local adaptation of populations to the local demand of their surrounding - often-unidentified - environmental conditions (Edelaar et al., 2011; Ovaskainen et al., 2011; Whitlock, 2008; Whitlock & Gilbert, 2012). They highlight how some traits involved with adaptive evolution at the population level can be used to identify the ecological pressures underlying natural selection. They revealed that some traits, but not all the traits involved with *A. majus* local adaptation, were under natural selection by the ecological conditions changing with altitude that are tightly linked to the climatic envelope of populations.

### Support for the local adaptation scenario

Our results showed that quantitative genetic divergence was higher amongst *A. majus* populations for ten of the thirteen traits under study than what could be explained by neutral evolutionary divergence. They imply that local adaptation has shaped the phenotypic diversity of *A. majus* populations across their geographic range. Local adaptation is detected in the majority of *Q*_ST_-*F*_ST_ comparisons (T. Leinonen, O’Hara, Cano, & Merila, 2008). Our finding is robust against a range of neutral evolution scenarios for these traits that were extrapolated from the distribution of *F*_ST_ values based on multiple putatively neutral markers (Whitlock, 2008). Furthermore, our approach excludes the possibility that plasticity rather than divergent selection might have generated this signal of phenotypic divergence because we used a common garden experiment, and included trait heritability estimates in *Q*_ST_ calculations (Benoit Pujol et al., 2008; Spitze, 1993). Ten of the thirteen traits under study harboured the signature of divergent selection but two traits (the time to flowering and the number of the leaves) did not show departure from the baseline scenario of neutral evolution. One particular trait (germination date) was in fact more similar among populations than expected under neutrality. A scenario of stabilising selection is classically extrapolated in the case of similar results (Lamy, Plomion, Kremer, & Delzon, 2012) but we argue that trait homogenisation caused during the experiment by phenotypic plasticity might be another plausible explanation.

The most likely evolutionary scenario applying to *A. majus* populations requires invoking a history of local adaptation in a complex background of gene flow and reproductive isolation. Neutral genetic divergence was weak but significant amongst populations. This indicates that mechanisms such as genetic drift have likely shaped differentially the genetic background of *A. majus* populations at the scale of their global geographic range. This can be interpreted as the genetic signature of reproductive isolation which restricted at least partly gene exchanges (Pujol et al., 2017). The mountain landscape in the Pyrenees was found to have likely reinforced reproductive isolation between *A. majus* populations (Pujol et al., 2017). However at the scale of *A. majus* neighbour populations from a contact zone, no genome wide barrier to gene flow was found (Ringbauer, Kolesnikov, Field, & Barton, 2018). At such local scale, gene flow is very likely rapidly homogenising population neutral genetic variation. Our findings thereby imply that Pyrenees mountains, which constitute a heterogeneous landscape that promotes complex patterns of connectivity amongst plant populations (Alberto et al., 2010), were prone to generate local adaptation in *A. majus*.

### The ecological significance of local adaptation in *A. majus*

In the absence of environmental measures included in the *Q*_ST_-*F*_ST_ analysis, it is impossible to identify the potential environmental agents of local selection that shape the quantitative genetic variation of traits. The functions behind the traits that have diverged during local adaptation can nevertheless be used to discuss plausible evolutionary scenarios of natural selection. Our results imply the adaptive divergence of flowering traits (flowering date, number of flowers, inflorescence size) amongst *A. majus* populations. These traits are directly linked to the reproductive success of populations and thereby often under natural selection (Meagher, 1994; Ollerton, Winfree, & Tarrant, 2011; Van Kleunen, 2007). They have great ecological significance in wild populations. Environmental constraints on the amount of resources available to plants are known to constrain flowering (Meagher & Delph, 2001). Biotic interactions between plants and pollinators are also known to modulate selection on flowering traits (Elzinga et al., 2007; Schemske & Bradshaw, 1999). Climate and seasonality differences amongst populations can also impose selection pressures on the flowering phenology (Franks, Sim, & Weis, 2007). Furthermore, populations flowering at different periods are less likely to exchange genes. Local adaptation shaping the flowering phenology can therefore have a feedback effect on population evolutionary divergence because it can reinforce this divergence by enhancing reproductive isolation. One plausible scenario of local adaptation is that a combination of these environmental pressures acts in fact as a complex agent of divergent selection amongst *A. majus* populations.

Our results also imply that local adaptation has shaped the vegetative architecture of plants that is specific to each *A. majus* population. They show that the quantitative genetic variation of several phenotypic traits characterising the vegetative growth and development of plants (stem diameter and length, number of nodes and internode distance, number of branches) has diverged among populations as a result of local adaptation. It is difficult to identify the environmental pressures underlying this finding because several environmental parameters (vegetation cover, wind, disturbance, temperature, water availability, etc.) can affect these traits. Every combination of them might have differed and be at the origin of divergent selection pressures between *A. majus* populations. Divergence in the genetic variation underlying the shape and size of plants was already found at the level of *Antirrhinum* species but its adaptive significance was not tested for (Langlade et al., 2005). The fact that a physiological trait, the Specific Leaf Area, is also involved with the local adaptation of *A. majus* populations suggests that the resource use ecological strategy of plants differs amongst populations. This is because population differences in this trait often reflect evolutionary divergence in the relative investment of resources in rapid growth (B. Pujol et al., 2008; Wilson, Thompson, & Hodgson, 1999).

Results like ours on the germination date of *A. majus* populations - lower quantitative genetic differentiation than expected under a scenario of neutral evolution - are widely acknowledged to imply stabilizing selection amongst populations (Lamy et al., 2012). Stabilizing selection of the germination date is a surprising finding because the timing of germination is often found to have evolved in response to divergent selection between habitats and populations (Donohue et al., 2005; Kalisz, 1986; Pujol et al., 2002; Weinig, 2000). The absence of fitness trade-offs between local environments might participate to explain this lack of divergence (Silvertown, 1981). The classical hypothesis suggested by this result is that a set of selective agents, which would be constant amongst populations, might act on this trait and favour a similar timing of germination across the species geographic and ecological range. Here we argue that an alternative but not mutually exclusive biological explanation can be invoked. There is evidence for the genetic basis of the germination timing in *A. majus* (Foley & Fennimore, 1998). Our alternative explanation nevertheless implies that the germination timing of *A. majus* populations was shaped by phenotypic plasticity in our experiment. For instance, we observed different abundance periods amongst *A. majus* populations in the wild. It is likely that phenotypic differences between wild populations that are not found in common garden conditions reflect plastic rather than genetic differences (Benoit Pujol et al., 2008). Here, we argue that the phenotypic similarity of populations in a common garden experiment might reflect their similar adjustment by means of phenotypic plasticity to the similar environment.

### Adaptive evolution of *A. majus* populations along the altitudinal gradient

Our results imply that the quantitative genetic basis of three of the thirteen traits under study (number of flowers, inflorescence length and germination date) was shaped by divergent selection between populations from different altitudes. Neutral genetic differentiation increased with population altitudinal differences, which means that smaller populations, less genetically diverse, and/or more reproductively isolated might be found higher on mountains (Pujol et al., 2017). This increase was however very small, so that neutral evolution is unlikely to explain the increase in quantitative genetic divergence observed for these three traits when population altitudinal differences increased. This latter increase was indeed one order of magnitude stronger than the baseline level set by neutral divergence (Figure 3). The hypothesis that the altitudinal gradient in quantitative genetic differentiation that we detected for three traits could be explained by neutral evolution therefore received poor support from our data. Contrasted environmental conditions characterise different altitudes in the geographic range of *A. majus*. In particular, we emphasize here the clear differences in rainfall, temperature and atmospheric pressure (Figure S2, Supporting Information). Other non-measured or unidentified ecological aspects (e.g., pollinator abundance, neighbouring plant community, vegetation cover) are also likely to change with altitude. These environmental factors are therefore good candidate selective agents for the observed adaptive evolutionary divergence of *A. majus* populations with altitude.

Most studies on plant adaptation to altitude report the selection of shorter plants that are faster growers at higher altitudes (Körner, 1999). There is also evidence for adaptive changes in flowering phenology and leaf traits to higher altitude environments (Körner, 1999). This is because there is generally a shorter growth season at higher altitudes, and thereby selection for fast developing plants that reproduce before the season deteriorates (Körner, 1999). Our results did not support a similar scenario of selection at play in *A. majus*. In contrast with the usual findings, our results showed that the divergent selection of flower production and germination timing between different altitudes predominates over flowering phenology, vegetative growth, and leaf traits in *A. majus* populations. *A. majus* plants that germinate later were selected for at higher altitudes. This is likely because it allows plants to track the later arrival of suitable climatic conditions for growth at higher altitudes (Körner, 1999). *A. majus* plants with smaller inflorescences and a reduced number of flowers were also selected for at higher altitudes. This can be the by-product of selection for the reduction of the whole plant size (Gonzalo-Turpin & Hazard, 2009). Alternatively, trade-offs in the resource allocation towards vegetative reproduction might be favoured at higher altitudes at the expense of sexual reproduction (Pluess & Stöcklin, 2005). The latter hypothesis is less likely in *A. majus* because we did not detect an increase in vegetative growth, and its vegetative reproduction in the wild is not documented. Our finding implies that environmental conditions becoming more stressful at higher altitudes (Körner, 2007) selected for reduced flower production.

Detecting the quantitative genetic signature of divergent selection imposed by environments that change with altitude has implications for the research on plant adaptation to climate change (De Villemereuil et al.). Our results provide evidence for the adaptation of *A. majus* populations to altitude in the Pyrenees, which might imply that *A. majus* successfully evolved adaptations to climate differences. The range of climate conditions in these mountains is already changing and set to change even more in the near future as a result of climate change. Conditions at high altitudes are becoming more similar to conditions from lower altitudes (Urli et al., 2014). Although signatures of past adaptive evolution have sometimes been used to evaluate the ability of populations to adapt to climate change on the long run, we argue that past adaptation does not imply that populations still have the ability to adapt. These signatures are however useful to identify potential adaptive traits, as for example flower production and germination date in *A. majus*, which might play a key role in the adaptation to climatic differences. The presence of heritable variation for adaptive traits is a more direct predictor of the ability of populations to respond to selection in the absence of specific constraints (Charmantier, Garant, & Kruuk, 2014; Kruuk, Slate, & Wilson, 2008; Pujol et al., 2018). In our approach, we quantified significant heritable variation in *A. majus* for traits potentially involved with adaptation to altitude. Under the hypothesis that no mechanisms will impede the response to selection in the wild (Pujol et al., 2018), our results suggest that *A. majus* populations might have some potential to adapt to climate change.

### Conclusion

Our findings corroborate the utility of *Q*_ST_-*F*_ST_ approaches conducted in common garden experiments to explore whether adaptive evolutionary divergence is required to explain trait quantitative genetic differentiation amongst populations. It is widely acknowledged that such approach can be used to identify traits that might be involved with local adaptation across the geographic range of a species. Our findings also participate to confirm the utility of the emerging use of *Q*_ST_-*F*_ST_ comparisons as a tool identifying candidate traits involved with the adaptation to selective agents acting on multiple populations rather than unknown local features of the environment. Here, we disentangled *A. majus* local adaptation from its response to selection along altitudinal gradients. This is a pertinent tool to explore how selection by ecological conditions that are currently changing, including climate envelopes, have already shaped the quantitative genetic variation of adaptive traits in the past. Combined with population estimates of the evolutionary potential to respond to selection, such approach might prove useful for building potential evolutionary scenarios of future adaptive evolution.

## Acknowledgments

We thank Jessica Cote for helpful discussions on the methods and David Field for helpful comments on the manuscript. This project has received funding from the European Research Council (ERC) under the European Union’s horizon 2020 research and innovation program (grant agreement No ERC-CoG-2015-681484-ANGI) awarded to BP. This work was supported by funding from the French “Agence Nationale de la Recherche” (ANR-13-JSV7-0002 “CAPA”) to BP. This project was also supported by the ANR funded French Laboratory of Excellence projects “LABEX TULIP” and “LABEX CEBA” (ANR-10-LABX-41, ANR-10-LABX-25-01).

## Supplementary Information

**FIGURE S1.**
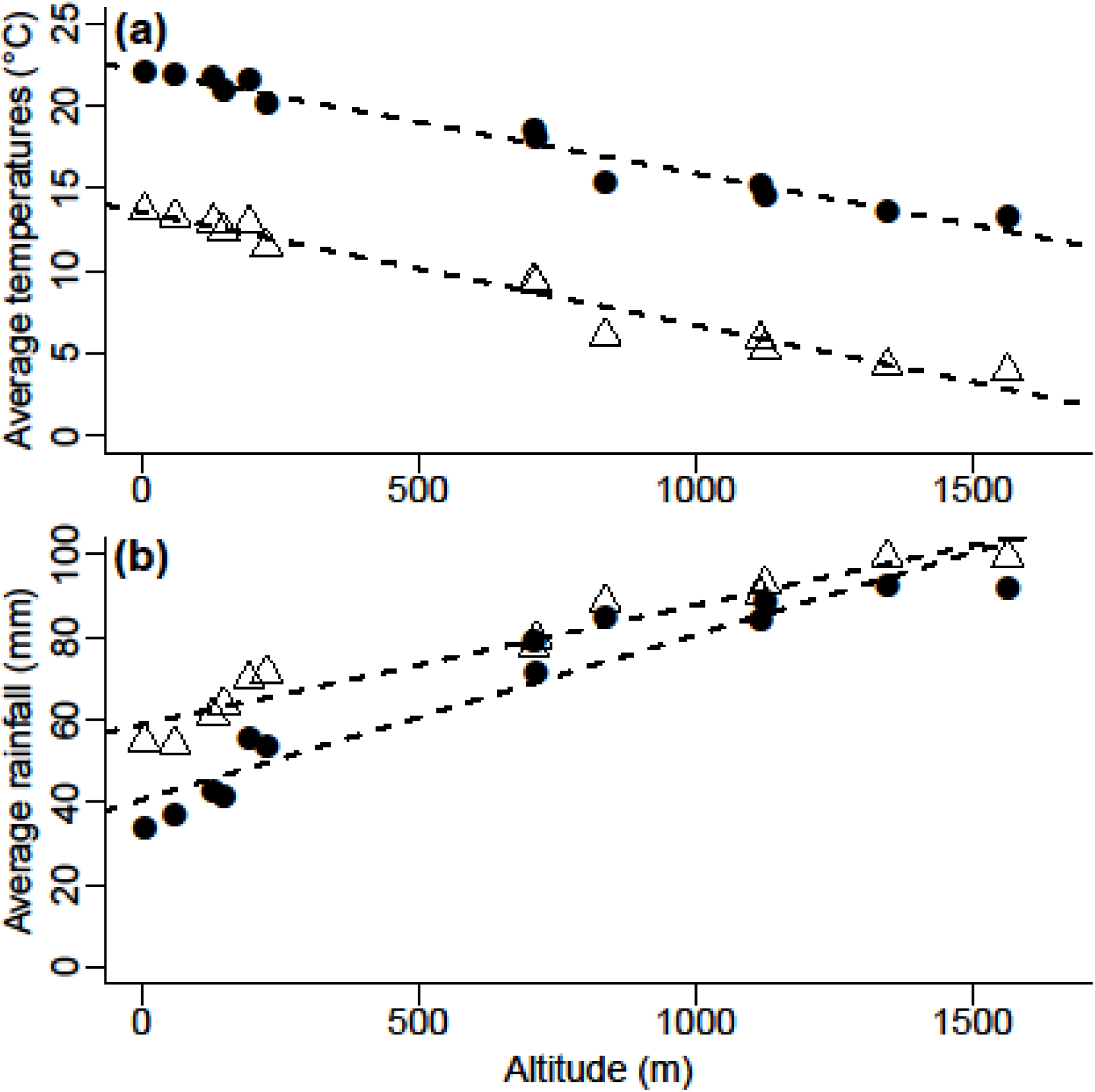
Spring and summer average temperatures and rainfall in the 13 populations. Population average temperature (a) and average rainfall (b) for the spring (triangles) and summer (dots) seasons as a function of altitude. Bioclimatic data was extracted from the *WorldClim* database (www.worldclim.org).

**FIGURE S1bis.**
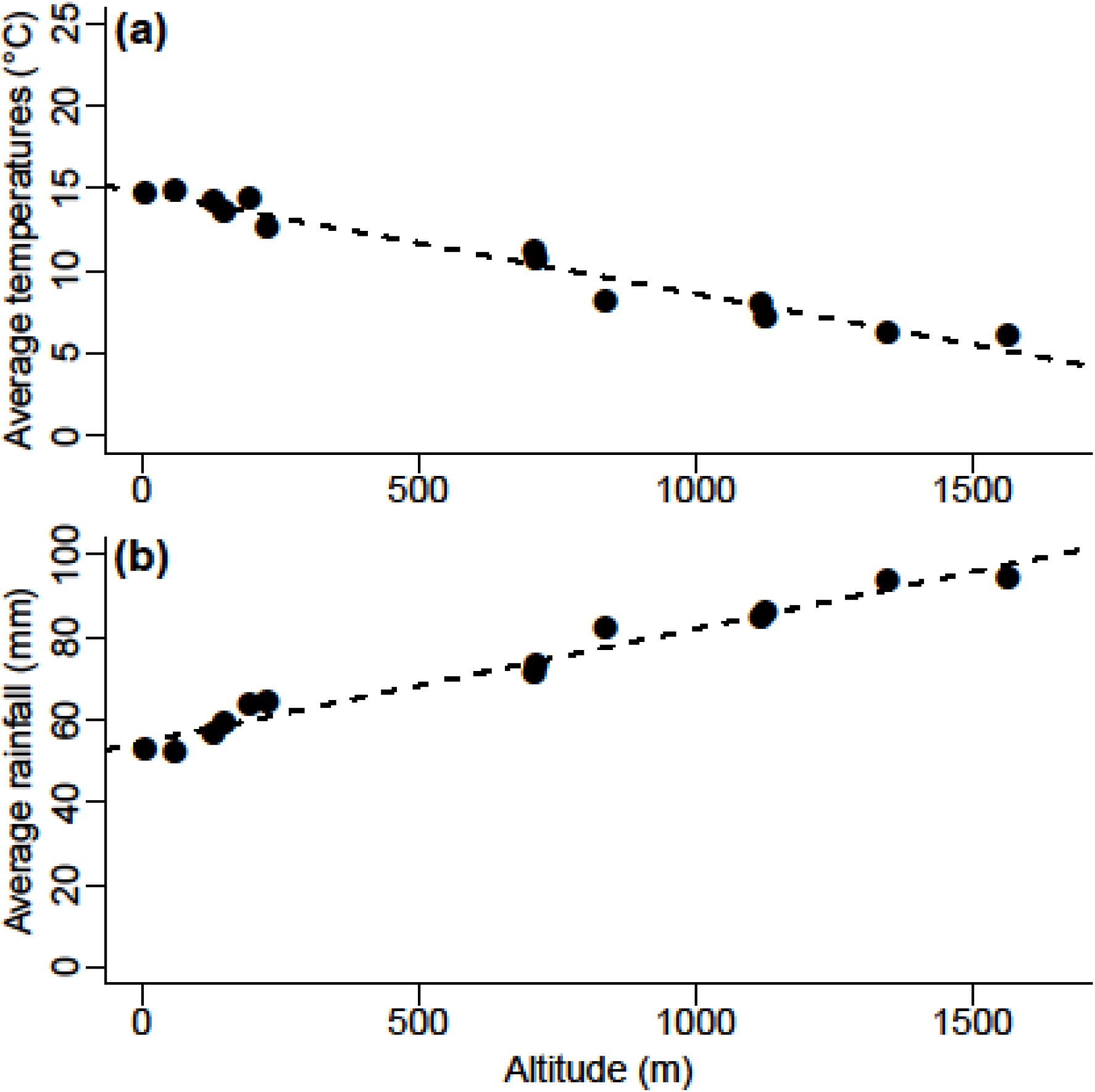
Annual average temperatures and rainfall in the 13 populations. Population average temperature (a) and average rainfall (b) as a function of altitude. Bioclimatic data was extracted from the *WorldClim* database (www.worldclim.org).

**FIGURE S2.**
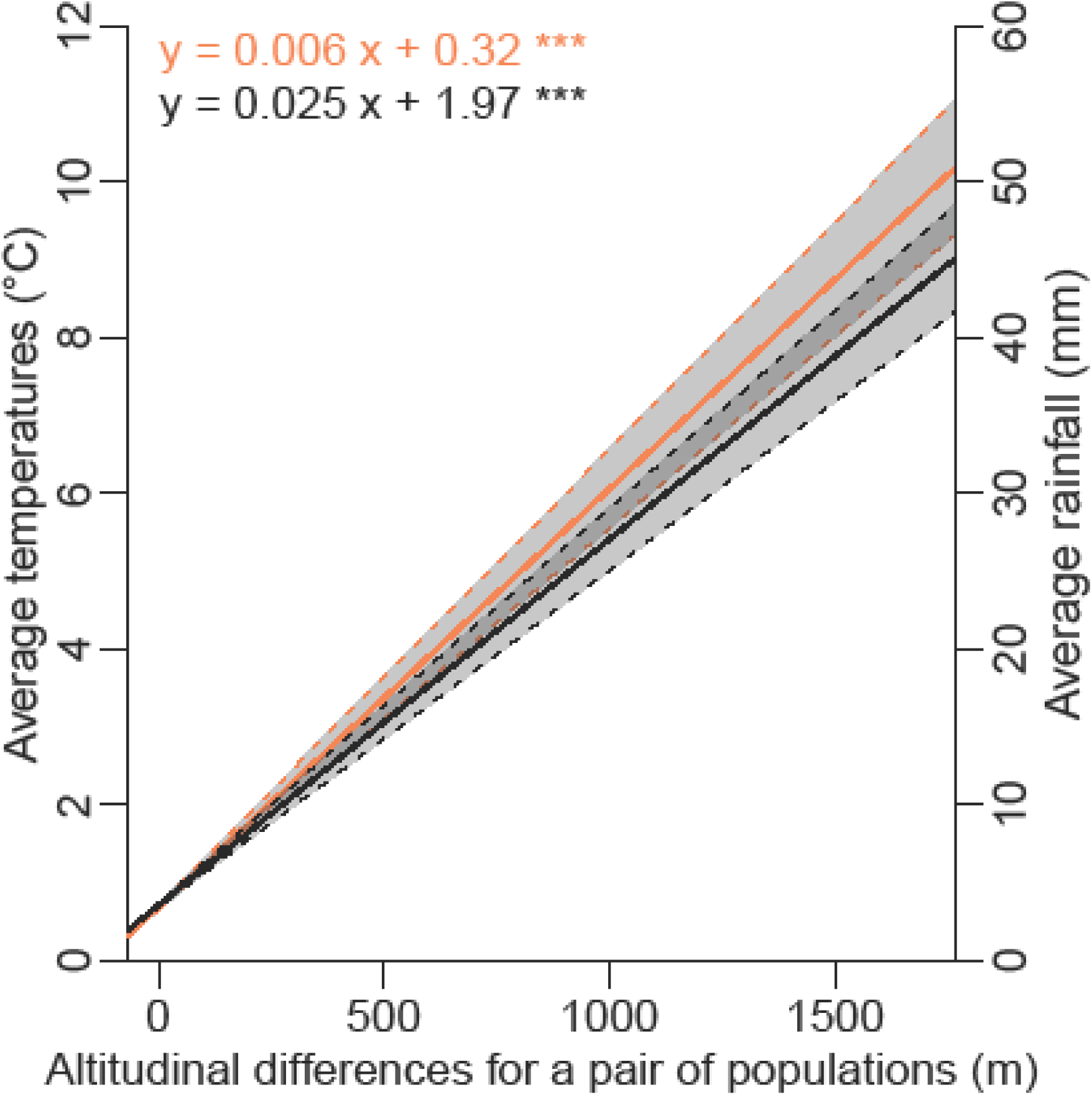
Linear regression lines of population pairwise average temperature (orange) and average rainfall (black) for a year with population pairwise differences in altitude (m). Regression equations and level of significance (*P*), obtained by Mantel test, are presented in figure and its 95% confidence interval is indicated by the shaded area. Bioclimatic data was extracted from the *WorldClim* database (www.worldclim.org).

**TABLE S1.**
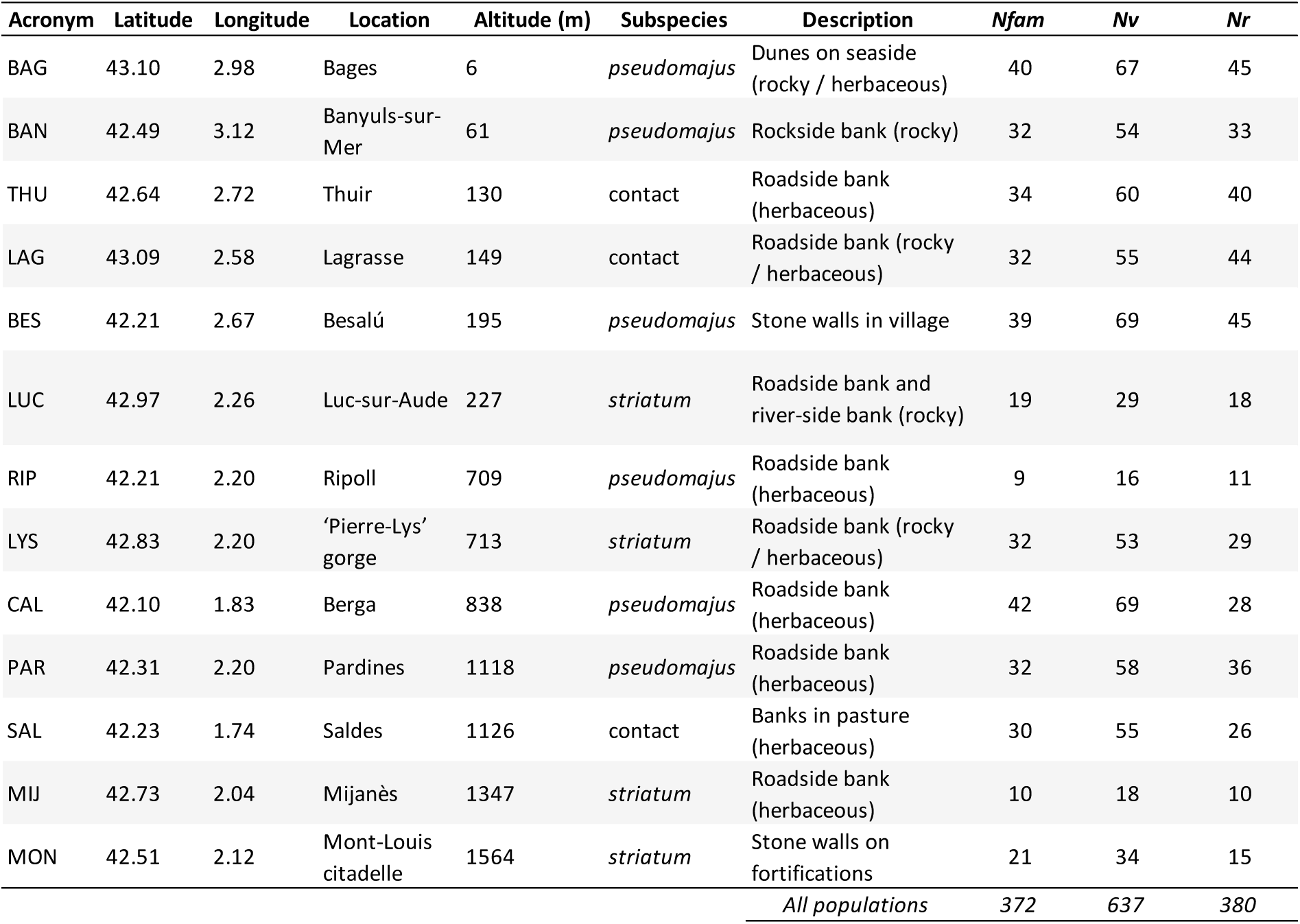
Description of the parental plant populations and characteristics of study system: number of families (*Nfam*), vegetative sampling size (*Nv*), reproductive sampling size (*Nr*).

**TABLE S2.** Summary of quantitative traits according to the sampling site. The table exhibits the average of phenotypic values of the common-garden experiment for all combined populations and different populations. All mean values are accompanied by their standard error.

| Phenotypic Traits | All<br>populations | BAG | BAN | THU | LAG | BES | LUC | RIP | LYS | CAL | PAR | SAL | MIJ | MON |
| --- | --- | --- | --- | --- | --- | --- | --- | --- | --- | --- | --- | --- | --- | --- |
| Germination date | 197.45<br>$\pm 0.45$ | 197.51<br>$\pm 1.38$ | 196.31<br>$\pm 1.37$ | 195.2<br>$\pm 1.32$ | 197.16<br>$\pm 1.62$ | 196.22<br>$\pm 1.18$ | 194.43<br>$\pm 1.48$ | 194.19<br>$\pm 2.55$ | 200.06<br>$\pm 1.7$ | 198.72<br>$\pm 1.74$ | 198.84<br>$\pm 1.43$ | 198.17<br>$\pm 1.48$ | 200.44<br>$\pm 2.94$ | 198.47<br>$\pm 1.83$ |
| Flowering date | 241.73<br>$\pm 0.66$ | 240.33<br>$\pm 1.87$ | 245.73<br>$\pm 2.47$ | 237.4<br>$\pm 1.24$ | 243.52<br>$\pm 2.86$ | 244.09<br>$\pm 1.81$ | 237.94<br>$\pm 2.15$ | 236.5<br>$\pm 1.34$ | 239.25<br>$\pm 2.55$ | 238.64<br>$\pm 1.3$ | 243.83<br>$\pm 1.84$ | 250.04<br>$\pm 3.12$ | 240.9<br>$\pm 3.18$ | 236.4<br>$\pm 1.05$ |
| Time to flowering | 47.97<br>$\pm 0.48$ | 46.56<br>$\pm 1.21$ | 51<br>$\pm 2.26$ | 45.5<br>$\pm 0.86$ | 48.45<br>$\pm 1.96$ | 49.40<br>$\pm 1.14$ | 44.67<br>$\pm 1.18$ | 48.5<br>$\pm 1.34$ | 45.11<br>$\pm 1.28$ | 45.96<br>$\pm 0.83$ | 48.83<br>$\pm 1.66$ | 56.36<br>$\pm 2.52$ | 47<br>$\pm 1.11$ | 43.8<br>$\pm 1.3$ |
| Diameter (mm) | 3.73<br>$\pm 0.04$ | 3.59<br>$\pm 0.10$ | 4.14<br>$\pm 0.15$ | 4.03<br>$\pm 0.12$ | 3.59<br>$\pm 0.11$ | 3.65<br>$\pm 0.13$ | 3.42<br>$\pm 0.15$ | 3.98<br>$\pm 0.19$ | 3.37<br>$\pm 0.14$ | 3.85<br>$\pm 0.11$ | 3.71<br>$\pm 0.14$ | 3.82<br>$\pm 0.13$ | 3.38<br>$\pm 0.18$ | 3.76<br>$\pm 0.16$ |
| Vegetative Length (cm) | 26.83<br>$\pm 0.34$ | 23.4<br>$\pm 0.79$ | 35.74<br>$\pm 1.18$ | 30.63<br>$\pm 0.71$ | 24.32<br>$\pm 0.89$ | 28.97<br>$\pm 1.18$ | 21.66<br>$\pm 0.85$ | 30.91<br>$\pm 2.1$ | 21.68<br>$\pm 0.88$ | 27.69<br>$\pm 1.02$ | 28.89<br>$\pm 1.28$ | 27.01<br>$\pm 1.01$ | 22<br>$\pm 1.52$ | 19.95<br>$\pm 0.73$ |
| Nodes | 13.21<br>$\pm 0.16$ | 12.46<br>$\pm 0.41$ | 15.07<br>$\pm 0.52$ | 13.33<br>$\pm 0.55$ | 11.07<br>$\pm 0.37$ | 14.67<br>$\pm 0.52$ | 12.69<br>$\pm 0.50$ | 16.38<br>$\pm 1.02$ | 11.62<br>$\pm 0.43$ | 13.7<br>$\pm 0.42$ | 12.95<br>$\pm 0.41$ | 15.89<br>$\pm 0.65$ | 10.61<br>$\pm 0.65$ | 10<br>$\pm 0.41$ |
| Leaves | 124.72<br>$\pm 2.12$ | 133.55<br>$\pm 6.35$ | 134.28<br>$\pm 7.73$ | 122.82<br>$\pm 6.5$ | 113.82<br>$\pm 5.74$ | 143.19<br>$\pm 8.6$ | 123.41<br>$\pm 11.29$ | 141<br>$\pm 12.56$ | 107.42<br>$\pm 5.67$ | 122.23<br>$\pm 4.89$ | 109.95<br>$\pm 6.54$ | 151.36<br>$\pm 7.47$ | 95.72<br>$\pm 10.7$ | 98.5<br>$\pm 5.82$ |
| Branches | 16.77<br>$\pm 0.29$ | 16.97<br>$\pm 0.75$ | 19.94<br>$\pm 1.21$ | 16.9<br>$\pm 0.73$ | 15.02<br>$\pm 0.78$ | 19.71<br>$\pm 1.19$ | 16.24<br>$\pm 1.76$ | 19.5<br>$\pm 1.8$ | 14.11<br>$\pm 0.77$ | 15.94<br>$\pm 0.77$ | 15.84<br>$\pm 0.9$ | 19.27<br>$\pm 0.88$ | 13.22<br>$\pm 1.55$ | 12.35<br>$\pm 0.67$ |
| Total Length (cm) | 45.18<br>$\pm 0.68$ | 43.74<br>$\pm 2.19$ | 54.49<br>$\pm 2.02$ | 52.29<br>$\pm 1.57$ | 43.84<br>$\pm 1.72$ | 45.94<br>$\pm 1.95$ | 35.03<br>$\pm 1.65$ | 48.49<br>$\pm 3.59$ | 38.96<br>$\pm 2.22$ | 49<br>$\pm 2.46$ | 51.14<br>$\pm 1.83$ | 41.28<br>$\pm 1.84$ | 36.66<br>$\pm 1.99$ | 25.18<br>$\pm 1.47$ |
| Inflorescence length (cm) | 16.82<br>$\pm 0.51$ | 20.00<br>$\pm 1.77$ | 16.42<br>$\pm 1.68$ | 21.92<br>$\pm 1.6$ | 18.58<br>$\pm 1.47$ | 15.92<br>$\pm 1.36$ | 11.83<br>$\pm 1.57$ | 15.15<br>$\pm 2.87$ | 15.03<br>$\pm 1.95$ | 17.52<br>$\pm 1.89$ | 18.47<br>$\pm 1.51$ | 12.79<br>$\pm 1.47$ | 11.57<br>$\pm 2.12$ | 6.86<br>$\pm 1.1$ |
| Flowers | 15.95<br>$\pm 0.49$ | 18.09<br>$\pm 1.70$ | 15.48<br>$\pm 1.54$ | 18.44<br>$\pm 1.34$ | 16.75<br>$\pm 1.41$ | 17.3<br>$\pm 1.43$ | 14.78<br>$\pm 2.42$ | 14.55<br>$\pm 2.86$ | 12.55<br>$\pm 1.46$ | 15.75<br>$\pm 1.72$ | 18.77<br>$\pm 1.73$ | 13.96<br>$\pm 1.46$ | 10.9<br>$\pm 2.09$ | 6.36<br>$\pm 1.1$ |
| Internode Length (cm) | 2.09<br>$\pm 0.02$ | 1.95<br>$\pm 0.07$ | 2.43<br>$\pm 0.08$ | 2.43<br>$\pm 0.07$ | 2.26<br>$\pm 0.08$ | 2.01<br>$\pm 0.07$ | 1.76<br>$\pm 0.09$ | 1.9<br>$\pm 0.08$ | 1.92<br>$\pm 0.08$ | 2.06<br>$\pm 0.07$ | 2.24<br>$\pm 0.08$ | 1.75<br>$\pm 0.06$ | 2.11<br>$\pm 0.13$ | 2.07<br>$\pm 0.09$ |
| SLA (cm<sup>2</sup>/g) | 176.08<br>$\pm 1.96$ | 173.29<br>$\pm 4.22$ | 178.44<br>$\pm 5.16$ | 179.55<br>$\pm 6.09$ | 182.1<br>$\pm 10.37$ | 194.48<br>$\pm 5.75$ | 161.24<br>$\pm 5.94$ | 165.9<br>$\pm 10.9$ | 179.53<br>$\pm 8.04$ | 171.29<br>$\pm 4.93$ | 189.06<br>$\pm 6$ | 162.62<br>$\pm 6.58$ | 178.74<br>$\pm 14.3$ | 144.58<br>$\pm 5.93$ |

**TABLE S3.** Differentiation for phenotypic traits among the 13 populations of *A. majus* grown in our common-garden experiment. Values for the within-population genetic variance (*Vw*), among-population genetic variance (*Vb*), heritability (*h²*) and observed (Obs.) *Q*_ST_ are given. *F*_ST_ overall populations was 0.107 (*P* < 0.001). *P-*values indicate statistical differences between observed and expected values of *Q*_ST-_*F*_ST_ under a neutrality hypothesis. The 95% CI of *Q*_ST_ (*Q*_ST_ .CI) and the *P* values of the *Q*_ST-_*F*_ST_ contrast are based on 10^5^ bootstrap replicates, following Whitlock & Guillaume (2009). Significant values are indicated in bold. The degrees of freedom used in the bootstrapping procedures are 12 for the among-population component (*Vb*) and are given in this table for the within-population component (df.*Vw*).

| Traits | $V_w$ | df.$V_w$ | $V_b$ | $h^2$ | $Q_{ST} \text{ obs}$ | $Q_{ST} \text{ CI}$ | $P \text{ -right}$ | $P \text{ -left}$ |
| --- | --- | --- | --- | --- | --- | --- | --- | --- |
| Germination date | 23.0964 | 356 | 0 | 0.36 | 0 | NA | 1 | **0** |
| Flowering date | 5.3673 | 259 | 9.828 | 0.07 | 0.48 | 0.247-0.644 | **0** | 1 |
| Time to flowering | 16.2058 | 259 | 7.4606 | 0.40 | 0.19 | 0.077-0.312 | 0.039 | 0.961 |
| Diameter | 0.0559 | 359 | 0.035 | 0.12 | 0.24 | 0.102-0.383 | **0.003** | 0.997 |
| Vegetative Length | 1.8973 | 359 | 19.9007 | 0.07 | 0.84 | 0.660-0.912 | **0** | 1 |
| Nodes | 3.1459 | 359 | 3.285 | 0.47 | 0.34 | 0.160-0.514 | **0** | 1 |
| Leaves | 0 | 359 | 217.3899 | 0 | 1 | NA | NA | NA |
| Branches | 1.7905 | 356 | 5.0482 | 0.08 | 0.59 | 0.332-0.736 | **0** | 1 |
| Total Length | 15.5028 | 266 | 57.5775 | 0.23 | 0.65 | 0.403-0.787 | **0** | 1 |
| Inflorescence length | 10.5535 | 263 | 10.7657 | 0.23 | 0.34 | 0.155-0.507 | **0** | 1 |
| Flowers | 6.8182 | 263 | 5.8013 | 0.16 | 0.30 | 0.133-0.462 | **0** | 1 |
| Internode Length | 0.0641 | 359 | 0.0428 | 0.42 | 0.25 | 0.107-0.397 | **0** | 1 |
| SLA | 146.6081 | 359 | 108.8237 | 0.13 | 0.27 | 0.118-0.422 | **0** | 1 |

